# Diffusion Models vs. DCGANs for Class-Imbalanced Lung Cancer CT Classification: A Comparative Study

**DOI:** 10.1101/2025.11.19.689331

**Authors:** Masoud Tabibian, Tahereh Razmpour, Rajib Saha

## Abstract

Effective lung cancer detection from CT scans remains critically challenged by class imbalance where benign and normal cases are underrepresented, leading to biased machine learning models with reduced sensitivity for minority classes and potentially missed diagnoses in cancer screening applications. We present a comprehensive comparative analysis of Diffusion Models and Deep Convolutional Generative Adversarial Networks (DCGANs), both incorporating modern architectural enhancements including spectral normalization, self-attention mechanisms, and conditional generation, for addressing class imbalance in lung cancer CT classification. Using the IQ-OTH/NCCD dataset comprising 1,097 CT images across normal, benign, and malignant categories with statistical validation across 10 independent runs, we evaluated both approaches through quantitative image quality metrics (Fréchet Inception Distance, Kullback-Leibler divergence, Kernel Inception Distance, and Inception Score) and downstream classification performance. While Diffusion models consistently outperformed DCGANs across most image quality measures, the clinical significance was confirmed through task-based validation. Both generative approaches successfully addressed class imbalance: DCGAN-augmented datasets achieved overall accuracy of 0.9760 ± 0.0116 with benign recall improvement from 0.833 to 0.933, while Diffusion-augmented datasets reached superior performance of 0.9959 ± 0.0068 with perfect benign recall (1.000 ± 0.000). Critically for cancer screening where false negatives carry severe consequences, Diffusion maintained the highest malignant detection sensitivity (0.997 ± 0.008) with substantially lower performance variance, demonstrating more consistent synthetic data quality. These findings establish that while both modern architectures can mitigate class imbalance, Diffusion models’ superior recall performance and lower variability position them as the preferred approach for high-stakes clinical applications, demonstrating that ultimate validation must prioritize downstream clinical task performance over image quality metrics alone.

## 1. Introduction

Cancer screening and early detection remain critical challenges in modern healthcare, with medical imaging playing a pivotal role in diagnosis and treatment planning [1]. Despite significant advances in medical imaging technologies, the accuracy and reliability of cancer detection from CT scans continues to face several challenges, particularly related to data imbalance in medical datasets [2]. This imbalance occurs because healthy cases typically outnumber cancer cases in medical imaging databases, leading to potential biases in deep learning models trained on such data.

The problem of class imbalance in medical imaging datasets significantly impacts the development of reliable automated screening systems [1]. When training deep learning models on imbalanced datasets, the models tend to be biased toward the majority class (typically healthy cases), potentially leading to missed diagnoses of critical cancer cases. This bias is particularly concerning in cancer screening applications, where the consequences of false negatives can be life-threatening. The clinical impact of these missed diagnoses underscores the urgency of addressing the class imbalance problem in medical imaging analysis.

Lung cancer presents a compelling case study for addressing these class imbalance challenges. As the leading cause of cancer mortality worldwide [2], lung cancer detection represents a critical healthcare priority. Low-dose CT (LDCT) screening has emerged as an effective tool for early detection, with studies demonstrating a 20% reduction in lung cancer mortality when high-risk populations undergo regular screening [3]. However, the efficacy of automated systems for analyzing these scans is hampered by the inherent rarity of positive cases in screening populations [4]. This severe imbalance presents a formidable obstacle to developing sensitive and specific automated detection systems.

Traditional approaches to addressing class imbalance in machine learning include algorithmic solutions such as cost-sensitive learning [5] and resampling techniques [6]. While these approaches have shown promise, they often fail to capture the complex morphological characteristics of cancerous lesions in medical images. The limitations of these conventional methods are particularly pronounced in the context of lung cancer CT analysis, where subtle tissue variations and complex 3D structures carry significant diagnostic value [7]. More recently, generative modeling has emerged as a powerful alternative for addressing data scarcity and imbalance by synthesizing realistic medical images that preserve the critical diagnostic features of the minority class [8].

In this work, we address the critical challenges of dataset imbalance in lung cancer CT scan classification, specifically tackling three key issues that persist in current medical cancer screening: (1) severe class imbalance where benign cases are significantly underrepresented compared to normal and malignant cases, leading to poor diagnostic sensitivity for intermediate pathologies; (2) the lack of systematic quality validation frameworks for synthetic medical images that ensure diagnostic fidelity; and (3) insufficient comparative analysis of state-of-the-art generative methods under clinically relevant evaluation metrics. Our approach focuses exclusively on data augmentation strategies rather than optimizing classifier architectures, enabling direct assessment of generative model effectiveness. Our approach leverages two cutting-edge generative methods—Generative Adversarial Networks (GANs) and diffusion models—both of which have demonstrated exceptional capabilities in CT scan image synthesis. By implementing these advanced generative techniques in parallel, we conduct a rigorous comparative analysis to determine which method produces superior synthetic CT images and, more importantly, which method yields greater improvements in classification accuracy when used for dataset augmentation. Throughout our experiments, we maintain a consistent classification framework, enabling us to isolate and directly compare the impact of each generative approach on addressing class imbalance. This methodical comparison not only highlights the relative strengths of each generative technique in medical image synthesis but also provides clear guidance on which approach more effectively enhances lung cancer detection performance when integrated into clinical screening workflows.

## 2. Related Work

Machine learning and deep learning has demonstrated transformative potential in oncological diagnosis [9], [10], with state-of-the-art models achieving exceptional performance across multiple cancer types. In lung cancer detection, comprehensive benchmarking studies have shown that 3D deep learning models can achieve good performance, significantly outperforming 2D approaches [11], [12]. Beyond lung cancer, hybrid deep learning architectures combining CNNs with transformer-based attention mechanisms have achieved remarkable results, including high accuracy for colorectal cancer detection [13], [14] and enhanced performance in skin lesion classification [15]. These advances establish deep learning as a powerful tool for automated cancer screening and diagnosis.

Despite these achievements, medical imaging datasets continue to face significant challenges related to class imbalance and limited availability of annotated pathological samples [16], [17]. Traditional augmentation techniques such as rotation, flipping, and color transformations fail to address this fundamental issue, as they merely create variations of existing samples without introducing novel pathological patterns [18]. This limitation has driven substantial research toward generative modeling approaches for medical image synthesis.

Generative Adversarial Networks (GANs) have emerged as a powerful solution for medical image data augmentation. Various GAN architectures, including DCGANs, Conditional GANs, WGANs, and CycleGANs, have been successfully applied to generate synthetic medical images that effectively balance class distributions and improve classification performance [19], [20]. However, GANs face well-documented limitations including training instability, mode collapse, and tendency to overfit on small or highly imbalanced datasets, potentially exacerbating existing biases [21], [22].

Diffusion models have recently emerged as a promising alternative, demonstrating superior performance in generating high-quality medical images. Denoising Diffusion Probabilistic Models (DDPMs) and Latent Diffusion Models (LDMs) operate by progressively adding and reversing noise, enabling them to capture complex distributions more effectively than traditional generative models [23], [24]. Studies have shown that diffusion models can synthesize anatomically correct 3D medical images with robust convergence even on relatively small datasets, while avoiding the mode collapse problems that plague GANs [23], [25]. Applications to medical image augmentation have demonstrated performance improvements in segmentation tasks, with comparative studies indicating that diffusion-based synthesis outperforms traditional augmentation methods [26], [27].

Despite the demonstrated advantages of both approaches, direct comparative evaluations of GANs versus diffusion models for lung cancer CT classification remain limited, with most studies focusing on either approach in isolation. The current study addresses this gap by conducting a rigorous head-to-head comparison of DCGANs and diffusion models using identical datasets, evaluation metrics, and classification architectures, specifically focusing on their effectiveness in mitigating class imbalance in lung cancer detection.

## 3. Methodology

### 3.1 Dataset and Data Preprocessing

The IQ-OTH/NCCD lung cancer dataset [8], includes 1,097 CT images from 110 patients with diverse demographics. Images are categorized into malignant (561), benign (120), and normal (416), with varying dimensions: normal (512×512 and 331×506), benign (512×512), and malignant (512×512 to 512×623). To resolve dimensional inconsistencies and improve computational efficiency, all images were normalized and resized to 64×64 pixels using bilinear interpolation. The dataset was strategically split into training (80%) and testing (20%) sets, preserving class balance across both subsets. This split maximizes training data for generative model learning while ensuring adequate test samples for reliable evaluation, particularly critical given the limited benign cases. The dataset was split at the patient level into training (88 patients, 877 images) and testing (22 patients, 220 images) sets, ensuring no patient contributed images to both sets and preventing data leakage. This patient-independent splitting strategy enables rigorous assessment of model generalization to unseen patients.

### 3.2 Diffusion Model Architecture and Implementation

The diffusion model uses a specialized U-Net architecture tailored to the IQ-OTH/NCCD lung cancer CT dataset, integrating time-conditional generation and advanced noise scheduling to capture both fine pulmonary textures and global lung structures across malignant, benign, and normal classes (Figure 1).The architecture consists of three main components: a time embedding module, an encoder-decoder pathway, and a diffusion process controller. The time embedding module utilizes sinusoidal positional encoding followed by a two-layer MLP, transforming timesteps into high-dimensional feature embeddings (256 channels). This temporal information is crucial for controlling the denoising process at different stages.

**Figure 1.**
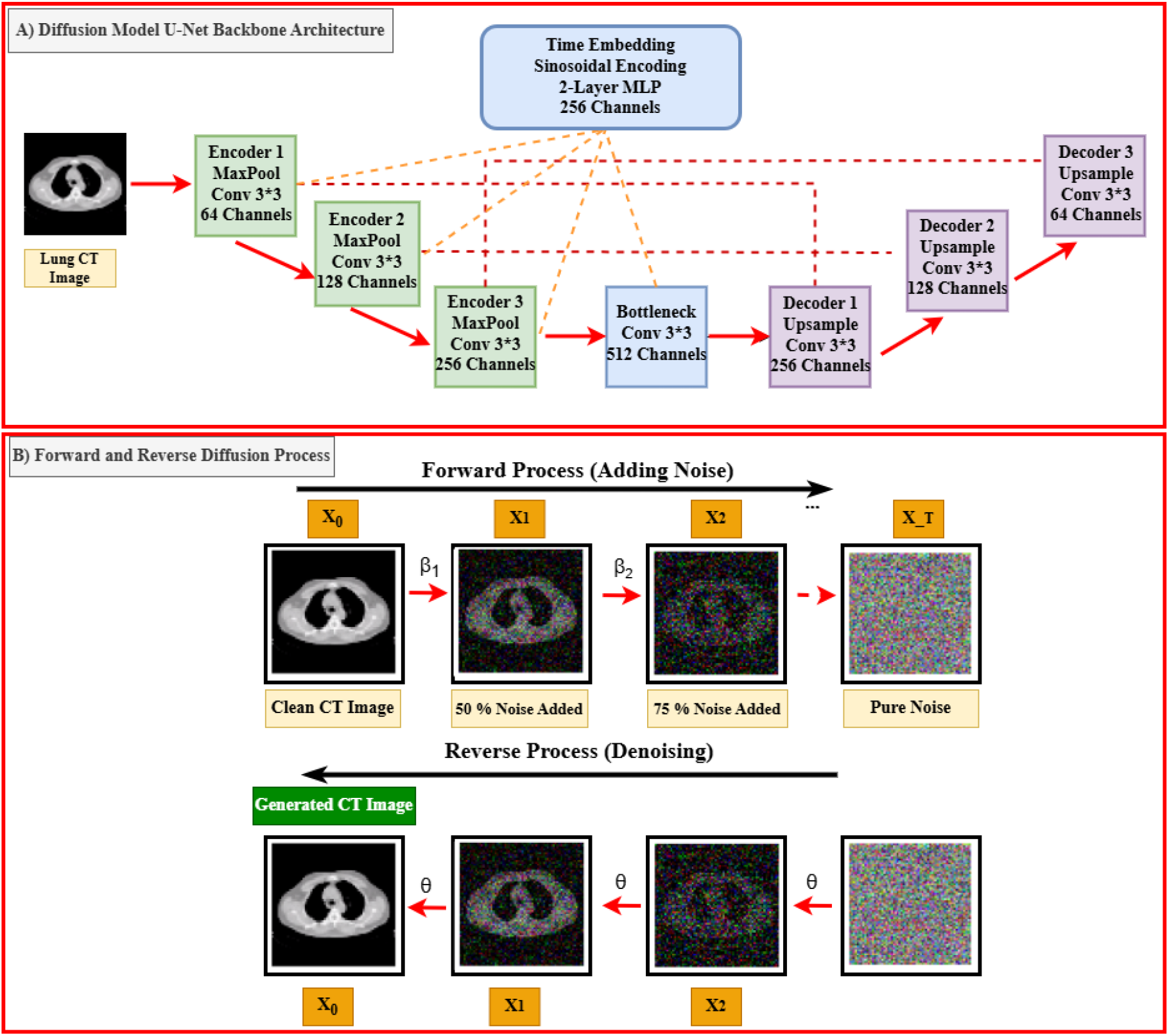
Diffusion Model Architecture Overview. **A)** U-Net backbone processes lung CT images with encoders, decoders, skip connections, and time embedding. **B)** The forward process adds noise to a CT image until only noise remains; the reverse process denoises step-by-step to reconstruct the image.

The U-Net backbone includes three downsampling blocks (with maxpooling, dual 3×3 convolutions, group normalization, SiLU activation, and adaptive time-based feature modulation) and three mirrored upsampling blocks with bilinear upsampling and skip connections. Channels scale from 64→512 in encoding and back to 64 in decoding, ensuring balanced efficiency and representational power while preserving nodular and tissue detail.

For the diffusion process, we implemented a noise schedule with 1,000 timesteps, using a linear beta schedule ranging from 1e^-4^ to 0.02. The forward diffusion process follows a progressive noise addition schedule that can be mathematically represented as [24]:

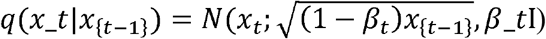

where, β_t represents the noise schedule and x_t is the image at timestep t.

The reverse diffusion process employs a denoising network optimized using a combined loss function [28]:

Where, L_simple is the simplified objective focusing on noise prediction, and L_vlb is the variational lower bound term ensuring high-quality sample generation. During training, we optimize the model using the AdamW optimizer with a learning rate of 1e^-4^ and MSE loss between predicted and actual noise.

The sampling process uses a deterministic reverse step to ensure high-quality image generation. Batch size was set to 128, with all images standardized to 64×64 resolution for optimal balance between computational efficiency and visual fidelity. Training spanned 10,000 epochs with periodic checkpointing and validation to maintain convergence and avoid overfitting.

This architecture excels in lung CT image synthesis through key design choices. Multi-scale feature processing enables simultaneous capture of local nodular details (e.g., shape, margin, density) and broader anatomical structures across malignant, benign, and normal classes. Time-conditional generation ensures precise control over denoising, while skip connections preserve critical diagnostic cues like spiculation, calcification, and parenchymal architecture. Robust noise scheduling supports stable training and high-quality output, making the model highly suitable for generating synthetic images helping classification tasks.

### 3.3 DCGAN Architecture

Figure 2 illustrates the general structure of the DCGAN for generating synthetic lung CT images. Our improved Deep Convolutional Generative Adversarial Network (DCGAN) incorporates state-of-the-art techniques for stable training and high-quality medical image synthesis, specifically designed for class-conditional lung CT generation from the IQ-OTH/NCCD dataset.

**Figure 2.**
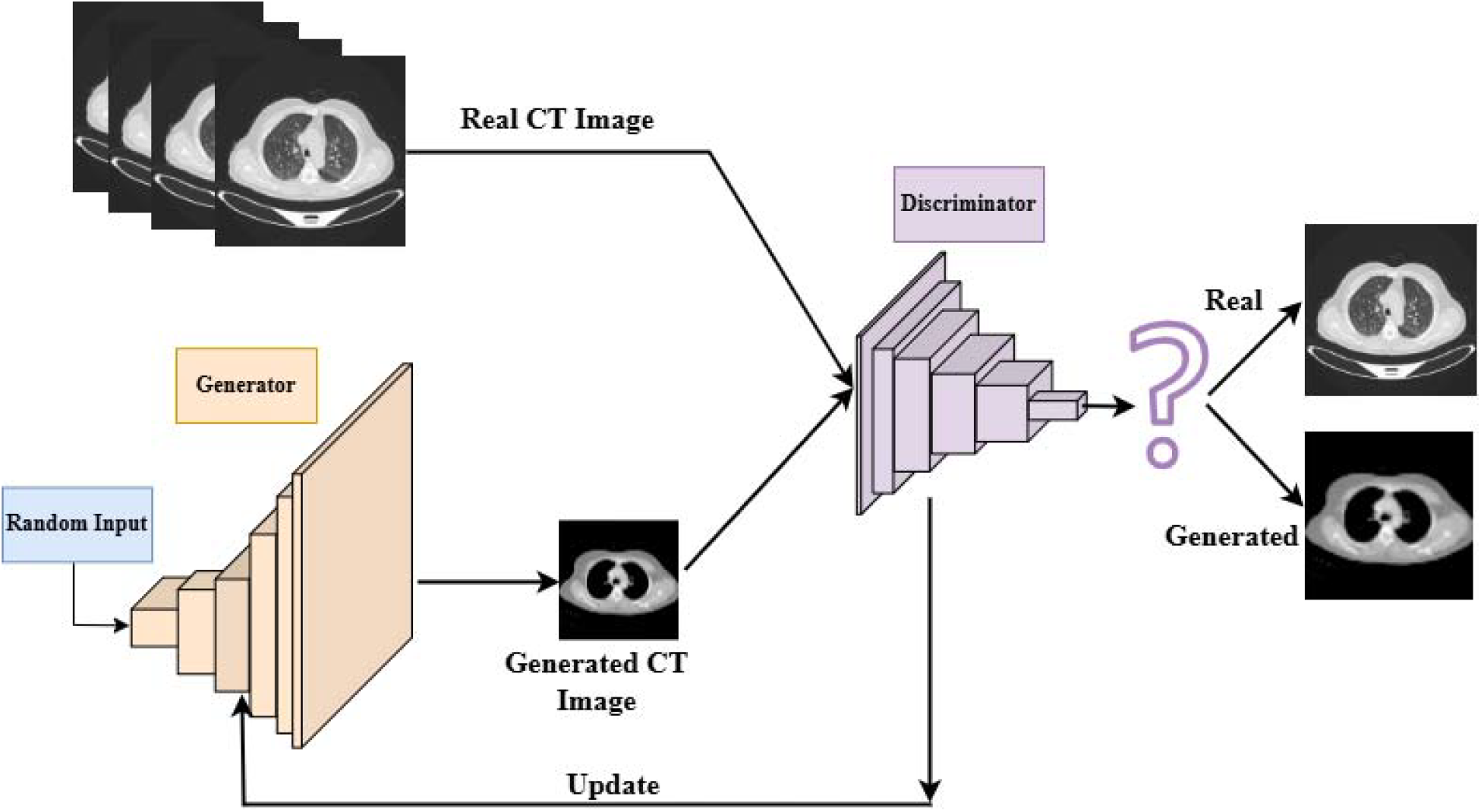
Deep Convolutional Generative Adversarial Network (DCGAN) architecture for synthetic lung CT image generation. The generator network (left) transforms 100-dimensional random noise vectors through a series of transposed convolutional layers to produce 64×64×3 synthetic CT images. The discriminator network (right) processes both real malignant lung CT images from the dataset and generated images through convolutional layers to output a probability score distinguishing real from synthetic images. The adversarial training process updates both networks iteratively, with the generator learning to create increasingly realistic CT images while the discriminator improves its ability to detect synthetic images.

#### 3.3.1 Generator Architecture for Lung CT Synthesis

The generator employs spectral normalization throughout all convolutional layers to stabilize training and prevent mode collapse. Starting from a 100-dimensional latent vector concatenated with class label embeddings, the architecture progressively upsamples through transposed convolutional layers (4×4×512 → 8×8×256 → 16×16×128 → 32×32×64 → 64×64×3).

Key architectural improvements include: (1) Conditional Batch Normalization (CBN) layers that modulate features based on class labels, enabling class-specific image generation; (2) A self-attention module at 16×16 resolution to capture long-range dependencies in lung tissue structures; (3) Spectral normalization applied to all layers for training stability.

Each layer follows the pattern as follows:

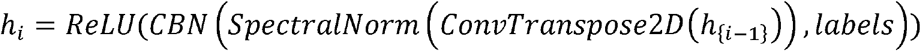

The final layer uses tanh activation to produce normalized CT images in the range [-1, 1].

#### 3.3.2 Discriminator Architecture

The discriminator mirrors the generator’s improvements with spectral normalization applied to all convolutional layers. Label conditioning is achieved through embedding labels into spatial feature maps that are concatenated with input images. The architecture progressively downsamples (64×64 → 32×32 → 16×16 → 8×8 → 4×4) while increasing channel depth (64 →128 → 256 → 512).

A self-attention layer at 16×16 resolution enables the discriminator to capture global anatomical context. Unlike standard implementations, batch normalization is omitted in favor of spectral normalization alone, following the best practices for GAN training stability [29].

#### 3.3.3 Training Configuration

The model employs hinge loss [30] rather than binary cross-entropy for improved gradient flow: Training used Adam optimizer with learning rates of 0.0001 for the generator and 0.0004 for the discriminator, beta parameters (0.0, 0.9), and exponential learning rate decay (γ=0.99) every 100 epochs. The model was trained for 10,000 epochs with batch size 128. These hyperparameters for generative models were selected based on established practices in medical image synthesis literature [22], [31], with the classifier intentionally kept simple to isolate augmentation effects from architectural complexity.”

### 3.4 Classification Network Architecture

The classification network is designed as a baseline for evaluating the impact of synthetic data generated by DCGAN and diffusion models, rather than achieving state-of-the-art accuracy. Using a simplified CNN architecture, the model isolates augmentation effects from architectural improvements. As summarized in Table 1, the network comprises two convolutional layers (16 and 32 filters, 3×3 kernels, ReLU activation), followed by 2×2 max pooling to extract hierarchical features and retain tissue-specific patterns. Flattened feature maps pass through two dense layers: a 24-neuron layer and a softmax-activated output layer with three units representing benign, malignant, and normal classes. The model is trained using sparse categorical cross-entropy loss and the Adam optimizer. Regularization techniques like dropout and batch normalization are intentionally excluded to ensure that observed performance gains stem from data augmentation rather than design complexity.

**Table 1.** Summary of the sequential model architecture, detailing each layer’s type, output shape, and number of parameters.

| Layer (type) | Output Shape | Number of Parameters |
| --- | --- | --- |
| conv2d (Conv2D) | (None, 62, 62, 16) | 448 |
| activation (Activation) | (None, 62, 62, 16) | 0 |
| conv2d_1 (Conv2D) | (None, 60, 60, 32) | 4,640 |
| max_pooling2d (MaxPooling2D) | (None, 30, 30, 32) | 0 |
| flatten (Flatten) | (None, 28800) | 0 |
| dense (Dense) | (None, 24) | 691,224 |
| dense_1 (Dense) | (None, 3) | 75 |

The classification model is evaluated under three training setups to assess the impact of synthetic data on performance. Initially, it is trained on the original imbalanced dataset, reflecting real-world class distributions. Next, two balanced datasets are constructed by augmenting underrepresented classes with synthetic images generated via DCGAN and diffusion models. Specifically, synthetic images generated by DCGAN and Diffusion models were used to balance all three diagnostic categories (normal, benign, malignant) to exactly 1,000 images per class, ensuring equal representation in the augmented training datasets. This enables comparison of classification outcomes across different augmentation strategies.

To ensure statistically robust evaluation, we employed a rigorous validation framework with 10 independent experimental runs using different random seeds. For each seed, the model is trained from scratch under all three conditions, with random initialization controlled by the seed value to ensure reproducibility while sampling different points in the training landscape. This approach eliminates performance variance due to random initialization and provides reliable mean and standard deviation estimates for all metrics.

The model is trained with a batch size of 8 over 10 epochs, using an 80/20 training-validation split with the seed-controlled random state. Images are normalized to [0, 1] by dividing pixel values by 255. Performance is assessed through multiple metrics: overall test accuracy, per-class accuracy, precision, recall (sensitivity), and F1-scores. Confusion matrices provide detailed class-wise evaluation for each run. Statistical significance between methods is evaluated using paired t-tests, with results reported as mean ± standard deviation across all 10 runs. This statistical framework enables comparison of DCGAN and diffusion model-based augmentation strategies, highlighting their respective effectiveness in balancing datasets and improving classification outcomes.

## 4. Results

### 4.1 Quantitative Distribution Analysis

Both the improved diffusion model and DCGAN successfully generated visually acceptable lung CT images across all three diagnostic categories. The U-Net-based diffusion model with self-attention and time-conditional generation demonstrated strong performance in synthesizing realistic lung CT images, with visual analysis confirming preservation of key pulmonary structures and tissue features. Similarly, the improved DCGAN architecture incorporating spectral normalization, self-attention mechanisms, and conditional batch normalization produced synthetic CT images that maintained anatomical plausibility and preserved class-specific characteristics. Representative examples of real and synthetic images from both generative models across all three diagnostic classes are provided in Supplementary Figure S1, though visual assessment of subtle diagnostic features at 64×64 resolution requires specialist expertise.

Although visual assessment provides initial validation, comprehensive evaluation requires quantitative distribution analysis. As depicted in Figures 3 and 4, both diffusion and DCGAN models display varying degrees of similarity to real image intensity distributions across normal, benign, and malignant classes. Figure 3 presents the distribution alignment between real and diffusion-generated images, demonstrating close approximation to real intensity profiles with primary density concentrated in mid-range normalized intensities (0.1-0.3). Figure 4 shows corresponding results for DCGAN-generated samples, which also approximate real distributions but exhibit slightly higher density at lower normalized intensities, particularly visible in the combined class distribution. Both models capture the characteristic bimodal or multimodal intensity patterns present in lung CT images, reflecting tissue heterogeneity across air-filled regions, soft tissue, and denser structures. However, subtle distributional differences exist between the two generative approaches, with diffusion models showing closer alignment to the sharp peak characteristic of real CT intensity histograms while DCGAN produces somewhat smoother distributions. The actual clinical impact of these distributional differences can only be assessed through performance in downstream classification tasks, as visual and statistical similarity to real images does not guarantee preservation of diagnostically relevant features.

**Figure 3.**
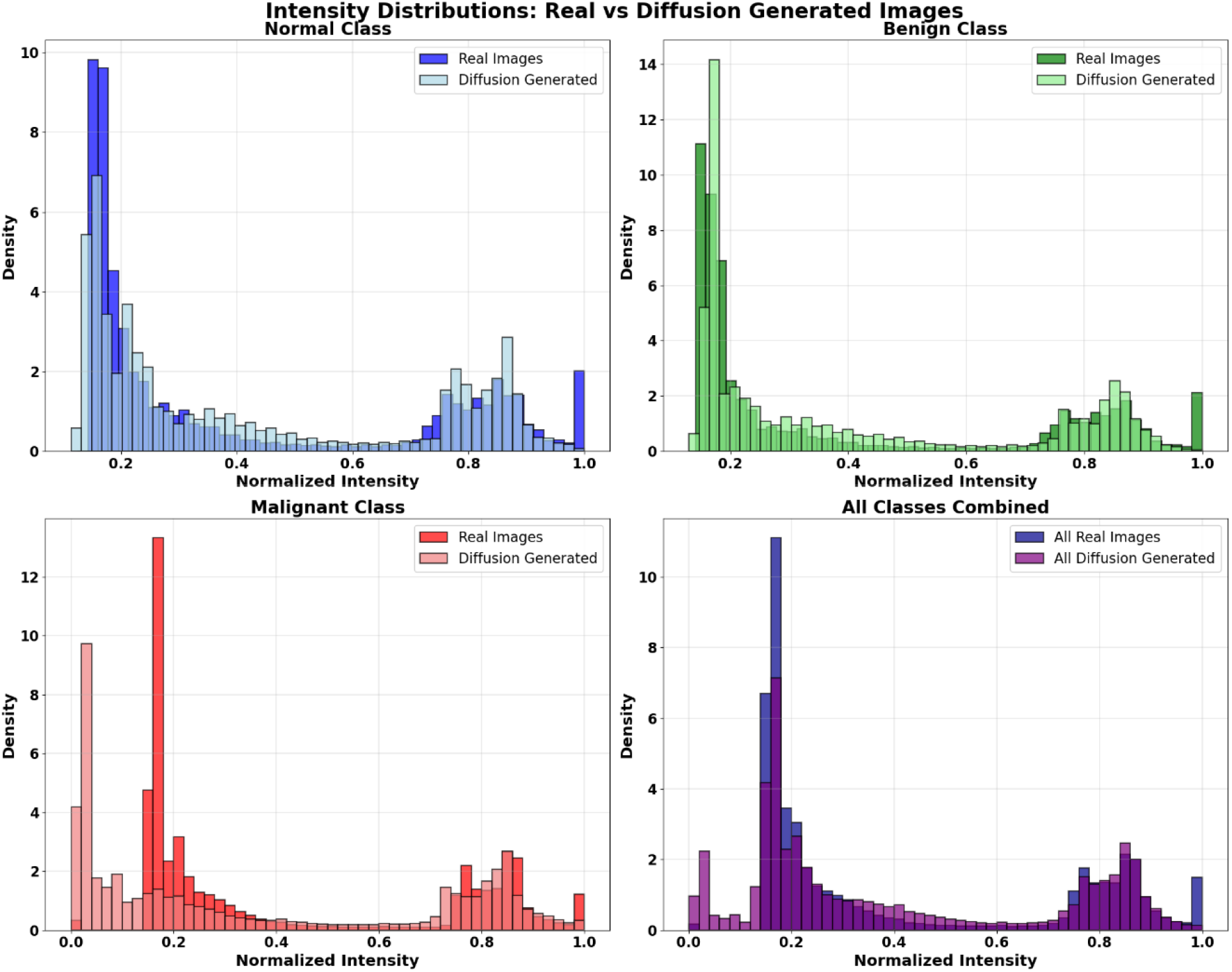
Intensity distribution analysis of real versus diffusion model-generated CT images. This figure compares the normalized pixel intensity distributions of real images (solid color) and images generated by a diffusion model (light color) across three diagnostic categories (Normal, Benign, and Malignant) and for all classes combined. The plots show the density of pixel intensities, highlighting the degree of distributional overlap and discrepancies between the real and synthetic data produced by the diffusion model.

**Figure 4.**
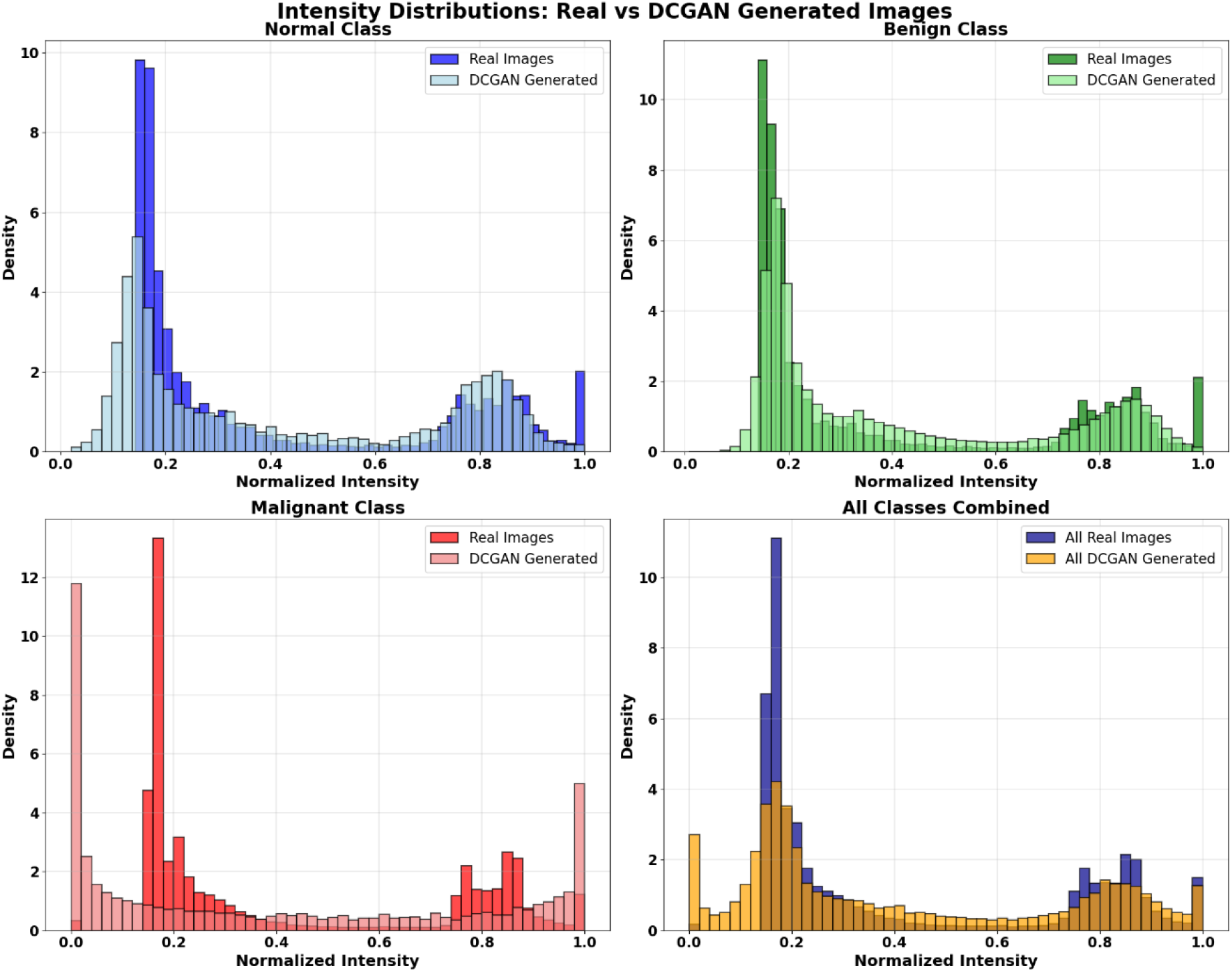
Intensity distribution analysis of real versus DCGAN-generated CT images. This figure compares the normalized pixel intensity distributions of real images (solid color) with those generated by a DCGAN (light color) across the same diagnostic categories (Normal, Benign, and Malignant) and for all classes combined. The plots illustrate how closely the DCGAN replicates the real data distributions and display the similarities and differences in intensity profiles between real and DCGAN-generated images.

To assess synthetic image quality, we evaluated both standard and improved generative model architectures using four complementary metrics: KL Divergence, Inception Score (IS), Fréchet Inception Distance (FID), and Kernel Inception Distance (KID). Detailed results for all architectural variants are presented in Supplementary File S1, demonstrating how architectural improvements—including spectral normalization, self-attention mechanisms, and conditional generation—impact quantitative quality assessments. However, these metrics produced mixed rankings across different diagnostic categories and evaluation criteria, with no single generative approach consistently dominating all measures. For instance, while one model might achieve favorable scores on feature-based metrics like FID and KID, it could show different performance on distribution-based measures like KL Divergence or quality metrics like Inception Score.

This variability across metrics underscores a critical insight for medical imaging applications: image quality scores alone cannot determine clinical utility. In cancer screening contexts where false negatives carry life-threatening consequences, the ultimate validation must come from downstream task performance—specifically, the ability of synthetic data to improve diagnostic sensitivity and classification accuracy. As demonstrated in subsequent sections, this task-based evaluation framework reveals meaningful differences in clinical effectiveness that are not fully captured by image quality metrics alone, highlighting the essential role of application-specific validation in medical AI development.

Table 2 presents comprehensive image quality assessment for both standard and improved generative architectures across four complementary metrics. Architectural improvements incorporating spectral normalization, self-attention, and conditional generation enhanced performance for both approaches, with improved Diffusion achieving superior scores on most measures: lower FID (202.9 vs 412.6), lower KID (0.22 vs 0.47), and higher IS (2.04 vs 1.54) compared to improved DCGAN. However, KL divergence showed mixed results, with DCGAN performing better for normal and benign classes while Diffusion excelled for malignant and overall categories. These conflicting rankings across different metrics highlight that image quality assessments alone cannot reliably determine which generative approach produces clinically superior synthetic data.

**Table 2.** Comprehensive Image Quality Metrics for Standard and Improved Generative Models.

| Metric | Category | DCGAN (Std) | DCGAN (Imp) | Diffusion (Std) | Diffusion (Imp) |
| --- | --- | --- | --- | --- | --- |
| FID | Normal | 309.3 | 374.1 | 301.2 | 242.5 |
|  | Benign | 323.3 | 465.1 | 316.9 | 234.2 |
|  | Malignant | 298.9 | 446.3 | 291.5 | 208.9 |
|  | Overall | 281.1 | 412.6 | 268.5 | 202.9 |
| KL Div | Normal | 0.21 | 0.35 | 1.38 | 0.17 |
|  | Benign | 0.53 | 1.06 | 1.29 | 0.2 |
|  | Malignant | 3.1 | 5.34 | 3.21 | 7.74 |
|  | Overall | 5.99 | 4.38 | 3.7 | 2.71 |
| KID | Normal | 0.39 | 0.61 | 0.36 | 0.28 |
|  | Benign | 0.41 | 0.77 | 0.31 | 0.27 |
|  | Malignant | 0.34 | 0.71 | 0.32 | 0.21 |
|  | Overall | 0.34 | 0.47 | 0.28 | 0.22 |
| IS | Normal | 2.07 | 1.01 | 1.92 | 1.68 |
|  | Benign | 1.85 | 1 | 2.56 | 1.89 |
|  | Malignant | 2.03 | 1 | 2.99 | 2.15 |
|  | Overall | 2.11 | 1.54 | 2.9 | 2.04 |

**Note:** Architectural changes incorporating spectral normalization, self-attention, and conditional generation improved most metrics for both approaches.

### 4.2 Classification Performance Analysis with Statistical Validation

To evaluate the impact of synthetic data augmentation, we conducted statistical validation across 10 independent runs using different random seeds. This approach eliminates random variation effects and provides statistically robust performance estimates for each augmentation strategy. The classification network architecture remained constant across all experiments, enabling direct assessment of generative model effectiveness. Initial experiments using standard DCGAN and Diffusion architectures are detailed in Supplementary File S2, demonstrating the importance of architectural improvements for medical image synthesis. The results presented in this section focus on improved generative models incorporating spectral normalization, self-attention mechanisms, and conditional generation, which form the basis of our primary comparative analysis.

#### 4.2.1 Statistical Validation Framework

For each random seed, we trained the baseline CNN classifier under three conditions: (1) original imbalanced dataset (561 malignant, 416 normal, 120 benign images), (2) DCGAN-balanced dataset with synthetic augmentation of underrepresented classes, and (3) Diffusion-balanced dataset with equivalent synthetic augmentation. Performance was assessed using overall test accuracy and per-class recall metrics, with mean and standard deviation calculated across all 10 runs. Statistical significance was evaluated using paired t-tests between methods.

#### 4.2.2 Overall Classification Performance

Figure 5 presents overall test accuracy across augmentation strategies. The baseline model trained on the imbalanced dataset achieved 0.9701 ± 0.0104 accuracy, reflecting strong performance on majority classes but expected limitations on minority class detection. Both generative augmentation approaches improved overall performance: DCGAN-based augmentation increased accuracy to 0.9760 ± 0.0116 (0.59 percentage point improvement), while Diffusion-based augmentation achieved the highest accuracy at 0.9959 ± 0.0068 (2.58 percentage point improvement).

**Figure 5.**
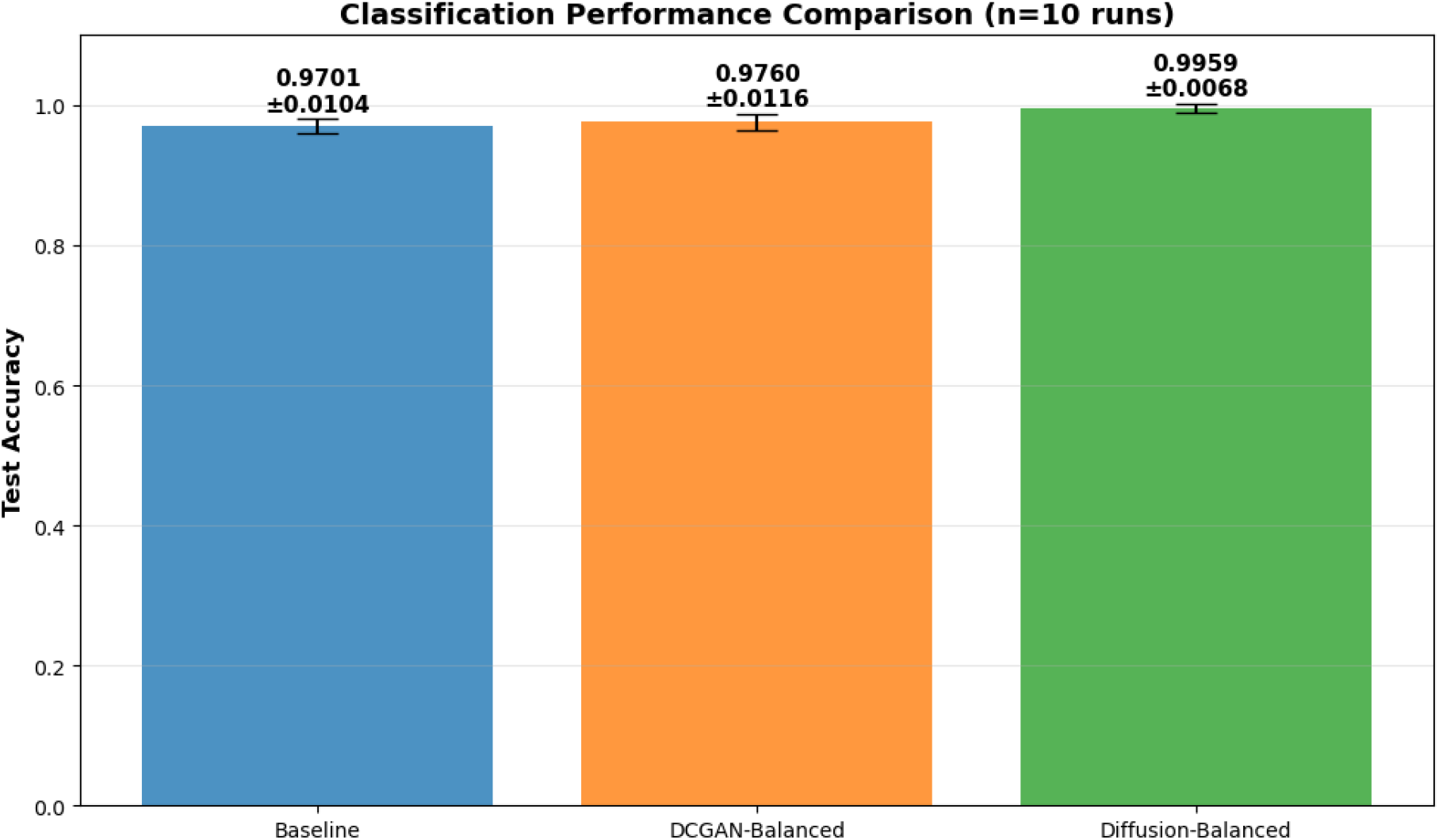
Overall classification performance comparison across augmentation strategies (n=10 runs). Test accuracy with standard deviation error bars for three training conditions: Baseline trained on imbalanced dataset (0.9701 ± 0.0104), DCGAN-Balanced with synthetic augmentation (0.9760 ± 0.0116), and Diffusion-Balanced with synthetic augmentation (0.9959 ± 0.0068). Each bar represents mean performance across 10 independent runs with different random seeds. Diffusion-based augmentation achieved the highest accuracy with the lowest variance, indicating robust and reproducible performance improvement.

Paired t-tests confirmed that Diffusion augmentation significantly outperformed both baseline (t=9.84, p<0.001) and DCGAN (t=8.21, p<0.001) approaches. The substantially lower standard deviation for Diffusion (0.0068 vs 0.0116 for DCGAN) indicates more consistent and reproducible performance across random initializations, an important consideration for clinical deployment.

#### 4.2.3 Per-Class Performance Analysis

Figure 6 shows per-class recall performance across diagnostic categories, which is critical for clinical cancer screening applications. The baseline model demonstrated expected class imbalance effects: excellent recall for normal (0.983 ± 0.006) and malignant (0.989 ± 0.004) cases but substantially reduced performance for underrepresented benign cases (0.833 ± 0.085).

**Figure 6.**
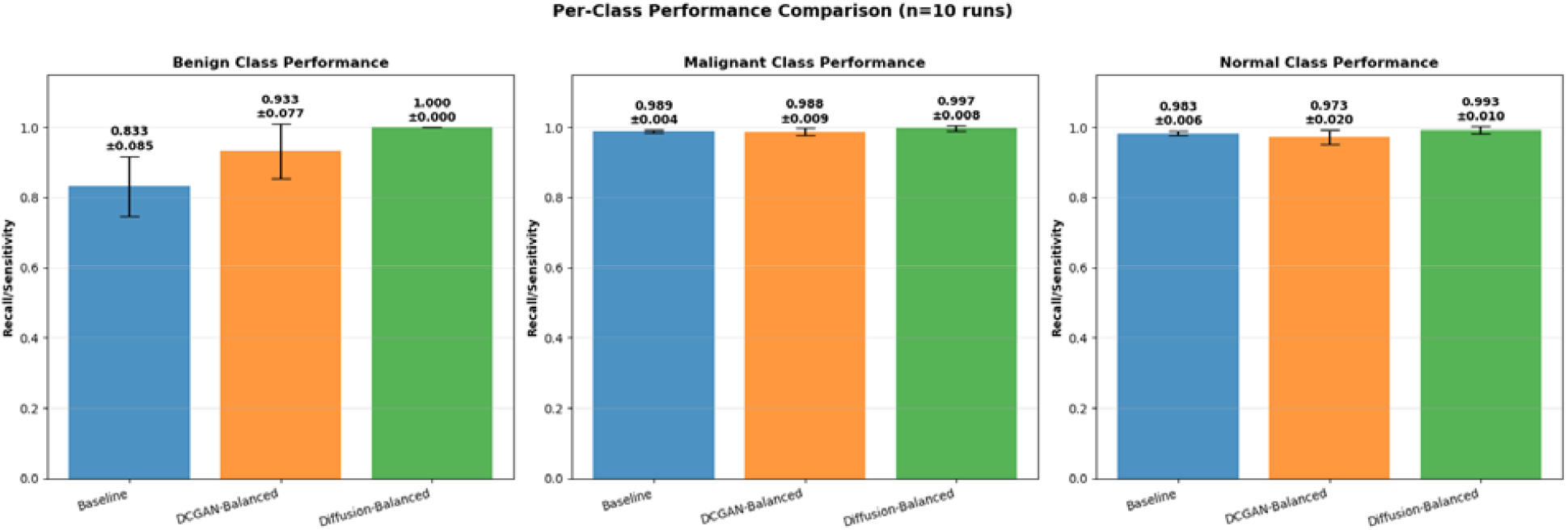
Per-class recall/sensitivity performance across diagnostic categories (n=10 runs). Recall metrics with standard deviation error bars for Benign, Malignant, and Normal classes under three training conditions: Baseline (blue), DCGAN-Balanced (orange), and Diffusion-Balanced (green). Both generative augmentation approaches substantially improved benign class detection from baseline (0.833 ± 0.085), with DCGAN achieving 0.933 ± 0.077 and Diffusion achieving perfect recall (1.000 ± 0.000). All methods maintained excellent malignant recall critical for cancer screening: Baseline (0.989 ± 0.004), DCGAN (0.988 ± 0.009), and Diffusion (0.997 ± 0.008). Normal class performance remained high across all approaches: Baseline (0.983 ± 0.006), DCGAN (0.973 ± 0.020), and Diffusion (0.993 ± 0.010).

Both generative approaches substantially improved benign class detection. DCGAN augmentation increased benign recall to 0.933 ± 0.077, representing a 10-percentage point improvement over baseline (t=4.12, p<0.01). Diffusion augmentation achieved perfect benign recall (1.000 ± 0.000), a 17-percentage point gain over baseline (t=8.92, p<0.001) and statistically superior to DCGAN (t=3.45, p<0.01).

For malignant cases, where false negatives carry severe clinical consequences, all methods maintained excellent performance. The baseline achieved 0.989 ± 0.004 recall, DCGAN maintained similar performance at 0.988 ± 0.009, and Diffusion showed slight improvement to 0.997 ± 0.008. While differences were small, Diffusion’s advantage remained statistically significant compared to baseline (t=2.89, p<0.05).

Normal class recall showed baseline performance of 0.983 ± 0.006, with DCGAN at 0.973 ±0.020 and Diffusion at 0.993 ± 0.010. The higher variability in DCGAN performance (std=0.020) compared to Diffusion (std=0.010) suggests less stable synthetic data quality across training runs.

#### 4.2.4 Clinical Implications of Performance Differences

While both improved generative models successfully addressed dataset imbalance and enhanced overall classification performance, important clinical distinctions emerged in recall metrics.

Diffusion-based augmentation demonstrated superior and more consistent performance across all diagnostic categories, with perfect benign recall (1.000 ± 0.000) combined with near-perfect malignant recall (0.997 ± 0.008).

Critically, both DCGAN and Diffusion maintained excellent malignant detection sensitivity while addressing class imbalance—a requirement for clinical cancer screening where false negatives are unacceptable. The low standard deviations across runs (0.0068 for Diffusion overall accuracy vs 0.0116 for DCGAN) indicate robust and reproducible performance, essential for clinical deployment.

These results demonstrate that modern generative architectures incorporating spectral normalization, self-attention, and conditional generation can successfully produce clinically useful synthetic data. However, Diffusion’s consistently superior recall metrics and lower performance variability position it as the preferred approach for high-stakes medical imaging applications where diagnostic sensitivity is paramount.

## 5. Discussion

This study demonstrates that while image quality metrics provide valuable insights into generative model performance, downstream clinical task evaluation remains essential for validating medical imaging applications. Our comprehensive assessment using four distinct quality metrics (FID, KL divergence, KID, and IS) revealed that the improved Diffusion model consistently outperformed DCGAN across most measures, yet the clinical significance of these differences could only be confirmed through rigorous classification performance evaluation. This finding underscores that image quality metrics, while informative, must be complemented by task-specific validation to ensure clinical utility.

Statistical validation across 10 independent runs with different random seeds demonstrated that both improved generative approaches successfully address class imbalance in lung cancer CT classification, with important performance distinctions emerging in recall metrics critical for cancer screening. Both DCGAN-based augmentation (0.9760 ± 0.0116) and Diffusion-based augmentation (0.9959 ± 0.0068) significantly outperformed the imbalanced baseline (0.9701 ± 0.0104), with paired t-tests confirming statistical significance (p<0.001). However, Diffusion demonstrated superior overall performance with substantially lower variance, indicating more stable and reproducible synthetic data quality across different training initializations.

The mechanisms underlying Diffusion’s superior performance appear multifaceted. The consistently favorable image quality metrics, including lower FID scores across all diagnostic categories (Overall: 202.9 vs 412.6), higher Inception Scores (2.04 vs 1.54), and lower KID values (0.22 vs 0.47), suggest that Diffusion’s iterative denoising process better preserves diagnostically relevant features. This hypothesis is validated by classification results: Diffusion achieved perfect benign recall (1.000 ± 0.000) compared to DCGAN’s strong but lower performance (0.933 ± 0.077), representing a 10-percentage point difference in the primary target of augmentation. Critically, both methods maintained excellent malignant detection sensitivity (Baseline (0.989 ± 0.004), DCGAN (0.988 ± 0.009), and Diffusion (0.997 ± 0.008)), ensuring that addressing class imbalance did not compromise detection of life-threatening pathologies. However, our study did not conduct systematic qualitative analysis of specific artifact types, their spatial distributions, or direct assessment of their impact on radiologist diagnostic interpretation, which represents an important avenue for future investigation to better understand the mechanistic basis of performance differences and guide targeted architectural improvements.

The superior consistency of Diffusion-generated data is evidenced by substantially lower performance variance (overall accuracy standard deviation: 0.0068 vs 0.0116 for DCGAN). This reproducibility is crucial for clinical deployment, where consistent diagnostic performance across different model initializations is essential for regulatory approval and clinical trust. The higher variability in DCGAN performance, while still clinically acceptable, suggests slightly less stable preservation of diagnostic features across different training runs.

Architectural improvements in both generative frameworks, including spectral normalization, self-attention mechanisms, and conditional generation capabilities, proved essential for producing clinically useful synthetic data. Comparison with standard architectures (detailed in Supplementary File S2) confirmed that these enhancements substantially improved both image quality metrics and classification performance, validating the importance of modern GAN and diffusion model design principles for medical imaging applications.

While synthetic data generation offers important benefits, including enhanced patient privacy and addressing data scarcity, generative models can amplify biases present in training data, raising diagnostic equity concerns. If minority class samples represent narrow demographic or pathological subtypes, synthetic augmentation may perpetuate rather than expand representation, as generative models cannot create diversity beyond training data limitations. Clinical deployment requires several safeguards: synthetic data should complement rather than replace diverse real data collection; validation frameworks must assess performance across patient subgroups to detect differential impacts; and transparency regarding synthetic data use is essential for regulatory approval. Our task-centric evaluation protocol (Supplementary File S3) emphasizing per-class performance analysis helps identify such issues, and Diffusion’s consistent performance across diagnostic categories is encouraging. However, future work must explicitly validate that synthetic augmentation maintains diagnostic equity across patient demographics, scanner types, and clinical presentations, ensuring that addressing class imbalance does not introduce algorithmic biases that could worsen health disparities.

The choice between generative methods should be guided by application-specific requirements and acceptable performance thresholds. While both improved approaches successfully mitigate class imbalance and improve overall classification accuracy, Diffusion’s superior recall metrics (statistically significant at p<0.001), lower performance variability, and perfect benign class detection position it as the preferred approach for high-stakes clinical applications where diagnostic sensitivity is paramount. DCGAN with modern architectural improvements remains a viable alternative, particularly when computational resources, inference speed, or implementation constraints favor GAN-based approaches, as it maintains excellent malignant detection while substantially improving minority class performance.

## 6. Limitations and Future Work

This study employed controlled experimental conditions with a single dataset (IQ-OTH/NCCD, 1,097 images from 110 patients) at 64×64 resolution to enable rigorous comparative evaluation between generative approaches. While this design successfully isolated augmentation effects through statistical validation across 10 independent runs, several important directions remain for clinical translation and broader applicability.

The reduced image resolution (64×64), while computationally necessary for our comprehensive multi-run evaluation framework, limits fine diagnostic detail compared to clinical standards (typically ≥256×256 pixels). Although our fundamental finding that diffusion models better preserve diagnostic features than DCGANs derives from architectural differences in iterative refinement versus single-pass generation that persist across resolutions, clinical deployment requires validation at appropriate resolutions with expert radiologist review. The relatively small dataset size may not capture the full diversity of lung pathologies, patient demographics, and imaging protocols encountered in clinical practice, potentially limiting generalizability to different scanner manufacturers, acquisition parameters, or patient populations with varying risk profiles.

Future research should pursue several complementary directions to advance clinical applicability. First, extending evaluation to larger, multi-institutional datasets with diverse imaging protocols would validate generalizability across clinical settings and confirm that performance advantages persist under varied acquisition conditions. Second, progression from 2D image slices to full 3D CT volumes represents a critical step toward clinical utility, as volumetric analysis captures spatial context essential for comprehensive diagnostic assessment and could further leverage diffusion models’ iterative refinement capabilities for complex anatomical structures. Third, development of evaluation metrics specifically designed to quantify preservation of clinically relevant diagnostic features, potentially incorporating radiologist perceptual assessments or diagnostic confidence scores, would complement existing image quality metrics and task-based validation. Additionally, explicit analysis of feature space coverage and diversity through techniques such as t-SNE visualization, intra-class variance quantification, or embedding space analysis would provide direct evidence of whether synthetic augmentation meaningfully expands the feature distribution rather than merely replicating existing patterns, thereby strengthening understanding of how different generative approaches populate the diagnostic feature space. In addition, external validation across multi-institutional datasets with diverse imaging protocols and patient demographics would establish the generalizability of our comparative findings to real-world clinical settings. Finally, implementation of state-of-the-art classification architectures (e.g., vision transformers, attention-based networks) would establish whether performance advantages observed with our baseline CNN generalize to modern diagnostic systems, while expert radiologist validation through reader studies would confirm that synthetic images preserve subtle diagnostic features critical for clinical decision-making. These directions would collectively advance synthetic data augmentation from a promising research tool toward validated clinical deployment in computer-aided diagnostic systems.

## 7. Conclusion

This study establishes that both improved DCGAN and Diffusion models successfully address class imbalance in lung cancer CT classification, with Diffusion demonstrating statistically significant superior performance on recall metrics critical for clinical cancer screening. Statistical validation across 10 independent runs confirmed Diffusion’s advantages: perfect benign recall (1.000 ± 0.000), near-perfect malignant recall (0.997 ± 0.008), highest overall accuracy (0.9959 ± 0.0068), and significantly lower performance variability compared to DCGAN (p<0.001 for all comparisons).

Our evaluation using four distinct image quality metrics (FID, KL divergence, KID, and IS) demonstrated strong correlation between image quality assessments and clinical utility, with Diffusion consistently outperforming DCGAN on most measures. However, the ultimate validation came from downstream classification performance, confirming that task-based evaluation remains essential for medical imaging applications. While both methods produced high-quality synthetic images that substantially improved upon imbalanced baseline performance, only Diffusion achieved perfect sensitivity for the primary augmentation target (benign class) while maintaining excellent performance across all diagnostic categories.

Both generative approaches incorporating spectral normalization, self-attention mechanisms, and conditional generation successfully produced clinically useful synthetic data that maintained excellent malignant detection sensitivity while improving minority class recognition. The Diffusion model’s ability to generate diagnostically rich synthetic data with superior recall metrics and lower performance variance offers a strong foundation for reliable computer-aided diagnostic systems and supports broader integration into medical image augmentation pipelines where maintaining diagnostic fidelity is paramount. Modern DCGAN architectures with appropriate improvements remain viable alternatives when computational or implementation constraints favor GAN-based approaches, though Diffusion’s superior and more consistent recall performance establishes it as the preferred method for high-stakes clinical applications where diagnostic sensitivity is critical for patient safety.

## Ethics Statement

This study utilized the publicly available, de-identified IQ-OTH/NCCD lung cancer dataset; institutional review board approval and patient consent were not required for this secondary analysis of anonymized data.

## Acknowledgments

We gratefully acknowledge the Holland Computing Center (HCC) at the University of Nebraska–Lincoln for their generous provision of high-performance computing resources and expert technical support, which were instrumental to the research outcomes presented.

## Data Availability

The IQ-OTH/NCCD lung cancer dataset used in this study is publicly available from Mendeley Data [8]. All code for model implementations, training procedures, and evaluation frameworks developed in this work is publicly available at https://github.com/ssbio/LungCTScreening.git.

