## Supplementary File S1 for "Diffusion Models vs. DCGANs for Class-Imbalanced Lung Cancer CT Classification: A Comparative Study"

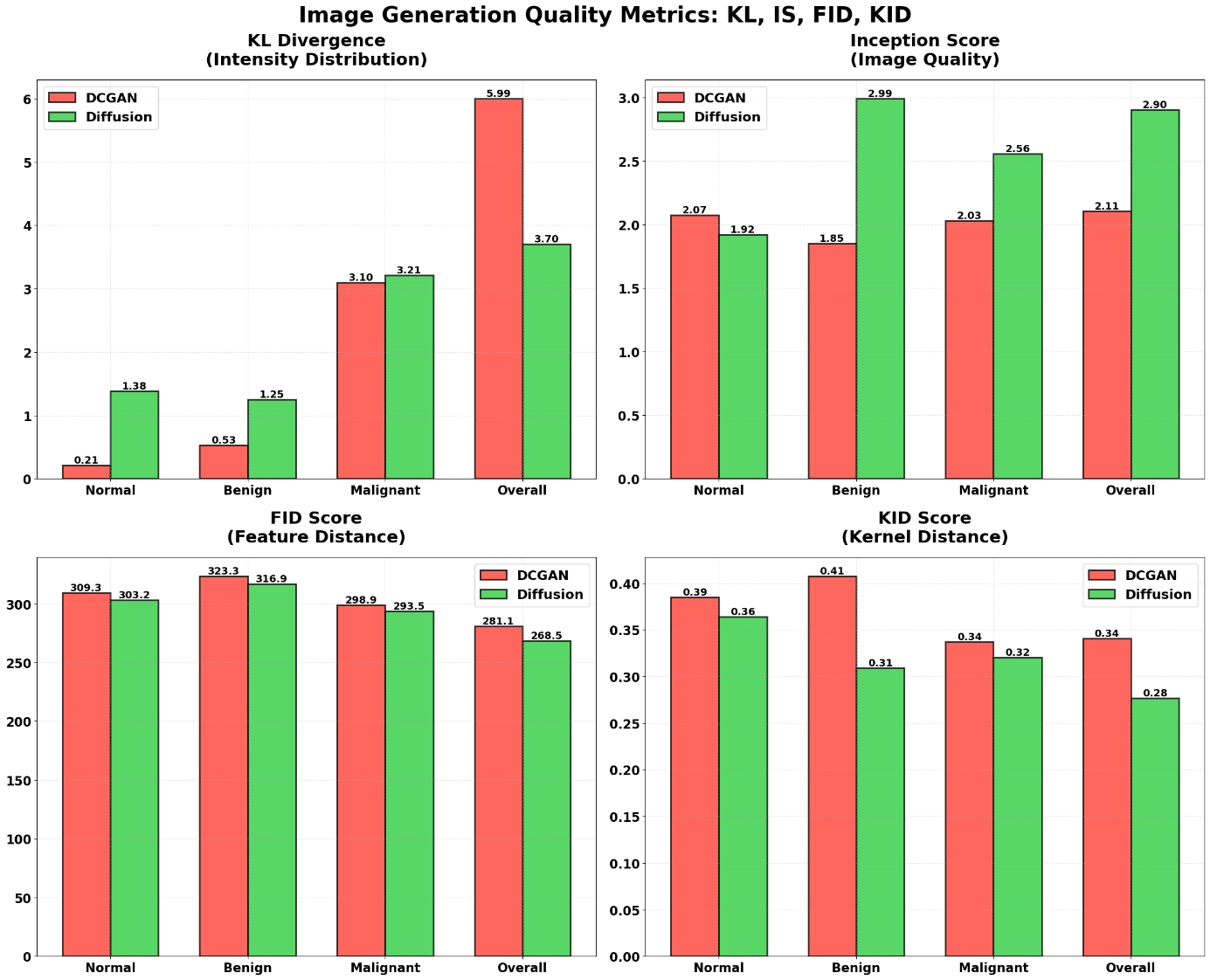

Quantitative assessment of synthetic lung CT images generated by basic generative models using four complementary metrics across diagnostic categories (Normal, Benign, Malignant, and Overall): (a) KL Divergence measuring intensity distribution similarity (lower indicates better alignment), (b) Inception Score evaluating image quality and diversity (higher indicates better quality), (c) Fréchet Inception Distance quantifying feature-level similarity (lower indicates closer match to real images), and (d) Kernel Inception Distance measuring distribution similarity in feature space (lower indicates better performance).

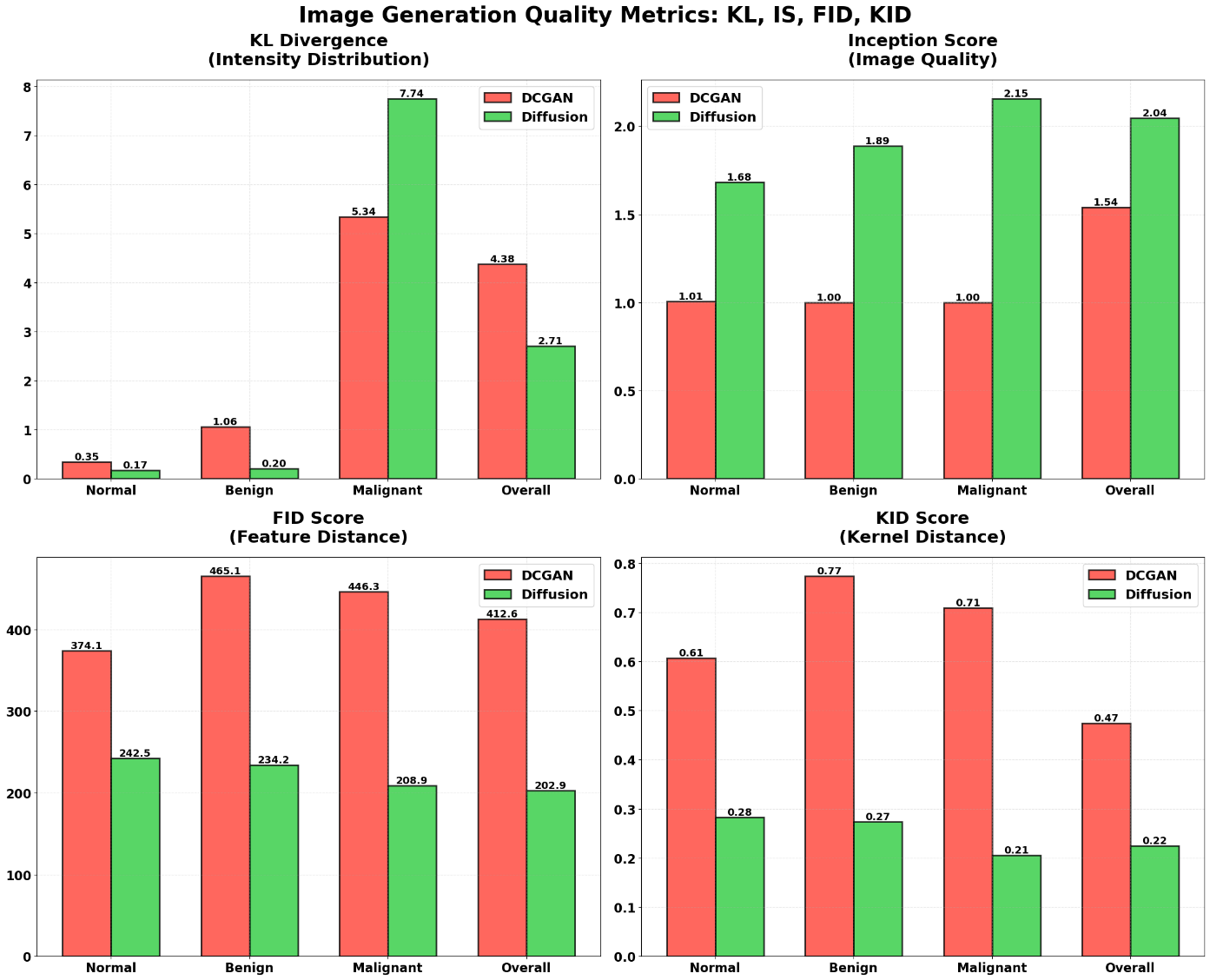

Quantitative assessment of synthetic lung CT images generated by improved generative models using four complementary metrics across diagnostic categories (Normal, Benign, Malignant, and Overall): (a) KL Divergence measuring intensity distribution similarity (lower indicates better alignment), (b) Inception Score evaluating image quality and diversity (higher indicates better quality), (c) Fréchet Inception Distance quantifying feature-level similarity (lower indicates closer match to real images), and (d) Kernel Inception Distance measuring distribution similarity in feature space (lower indicates better performance).
