## Supplementary File S2 for "Diffusion Models vs. DCGANs for Class-Imbalanced Lung Cancer CT Classification: A Comparative Study"

**Baseline Performance on Imbalanced Dataset**

To establish a performance baseline for subsequent comparisons, we first evaluated our classifier on the original imbalanced dataset. The original imbalanced dataset (561 malignant, 416 normal, 120 benign images) produced expected classification disparities. The model achieved 98% accuracy for normal and 97% for malignant cases, reflecting their dominance in the training set (Figure 1a), but struggled with benign cases, reaching only 67% accuracy and misclassifying 25% as normal. This performance pattern demonstrates how class imbalance leads to biased learning where the model becomes overly sensitive to majority classes while failing to capture the subtle features distinguishing minority classes. As shown in Figure 1b, closely aligned training and validation accuracy curves across epochs indicate stable learning without overfitting, confirming that the poor benign performance stems from insufficient training examples rather than model complexity issues. These baseline results establish the critical need for data augmentation, which we address through two distinct generative approaches in the following sections.

| (a) | (b) |
| --- | --- |
| 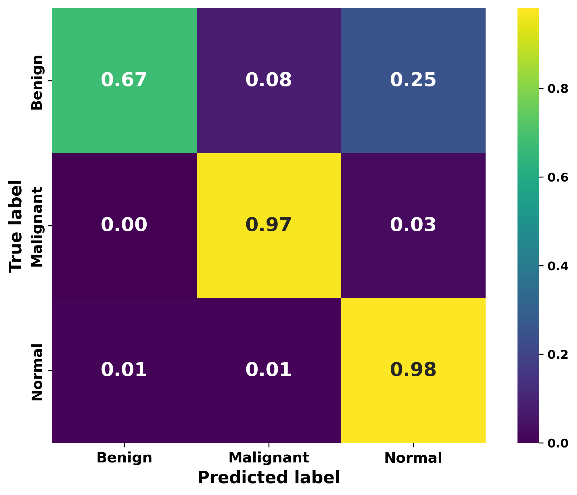 | 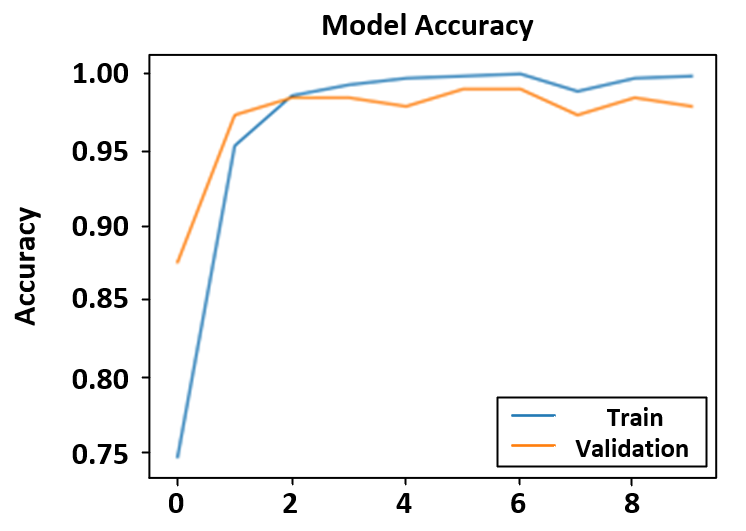 |
| **Figure 1. Confusion Matrix and Learning Curve for Baseline Classifier Applied on Imbalanced Dataset.**  **(a)**Confusion matrix showing classification performance on the original imbalanced IQ-OTH/NCCD dataset, with diagonal values representing correct classifications: Normal (0.98), Malignant (0.97), and Benign (0.67). The matrix reveals poor performance for benign cases, with 25% misclassified as normal, highlighting the impact of class imbalance. **(b)** Training and validation accuracy curves across 10 epochs demonstrate stable learning without overfitting, with both curves closely tracking each other and reaching high accuracy levels, indicating effective model generalization despite the dataset imbalance. | |

**Diffusion Model-Enhanced Performance**

Having established the baseline limitations, particularly the 67% accuracy for benign cases, we next evaluated whether diffusion-generated synthetic images could restore balanced classification performance. Balancing the dataset with diffusion-generated synthetic images significantly improved classification across all diagnostic categories. As shown in the confusion matrix (Figure 2a), the model achieved perfect accuracy (1.00) for normal cases and high accuracy (0.95) for malignant cases. The most notable gain was in benign case classification, which rose to 0.92—a 25-point increase over baseline—with only 8% misclassified as normal and none as malignant.

| (a) | (b) |
| --- | --- |
| 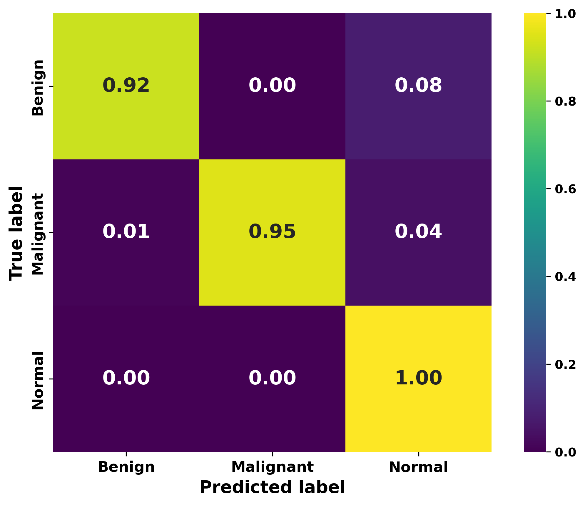 | 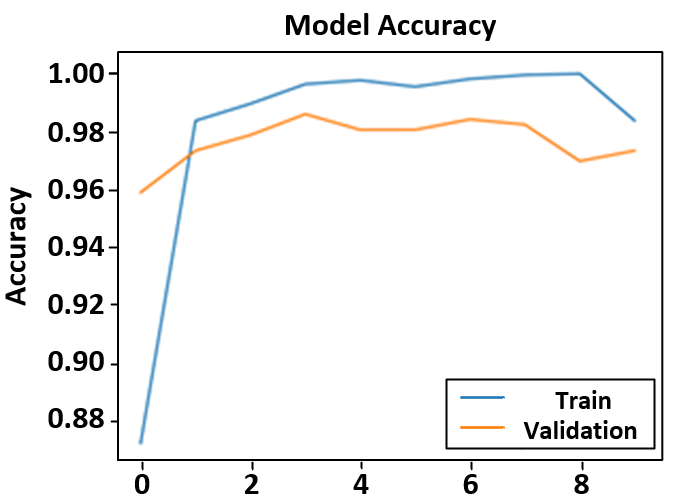 |
| **Figure 2. Confusion Matrix and Learning Curve for Classifier Applied on Diffusion-balanced Dataset.**  **(a)** Confusion matrix demonstrating substantially improved classification performance after diffusion model-based dataset balancing, with excellent diagonal accuracy values: Normal (1.00), Malignant (0.95), and Benign (0.92). The dramatic improvement in benign case classification (25 percentage points increase from baseline) with minimal off-diagonal misclassifications demonstrates the effectiveness of diffusion-generated synthetic data augmentation. **(b)** Training and validation accuracy curves showing rapid convergence to high performance levels and stable learning throughout 10 epochs, with both curves remaining consistently high and closely aligned, indicating excellent model generalization on the augmented dataset. | |

This performance boost came without compromising other class predictions, as shown by the low off-diagonal values in Figure 2a—highlighting the diagnostic strength of diffusion-generated images. The improvement stems from the model’s ability to produce diverse, high-quality synthetic samples that strengthened the benign class while preserving diagnostic features. The learning curves in Figure 2b confirm efficient learning and strong generalization, with training and validation accuracy remaining consistently high throughout epochs, indicating that the synthetic data enhances rather than confuses the learning process

**DCGAN-Enhanced Performance**

While the diffusion model demonstrated clear improvements across all classes, we next examined whether DCGAN-based augmentation could achieve similar benefits. In contrast to the diffusion model's success, the DCGAN-balanced dataset produced mixed classification results (Figure 3a). Normal case accuracy remained high at 0.99, and benign classification improved to 0.83—up 16 points from baseline. However, malignant case accuracy declined to 0.75, with 24% misclassified as normal, underscoring a key limitation of the DCGAN approach for this class.

| (a) | (b) |
| --- | --- |
| 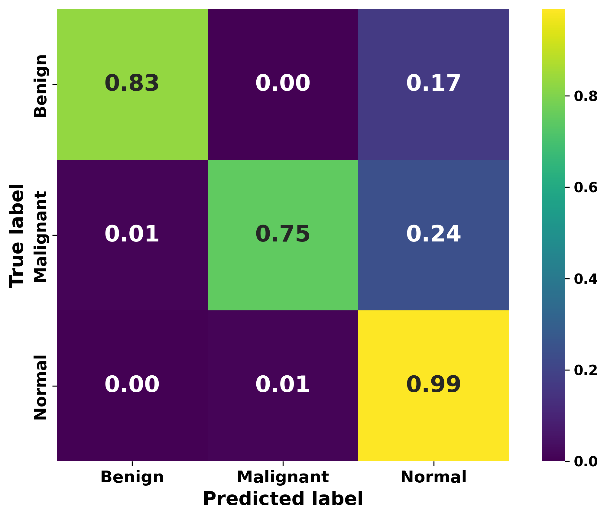 | 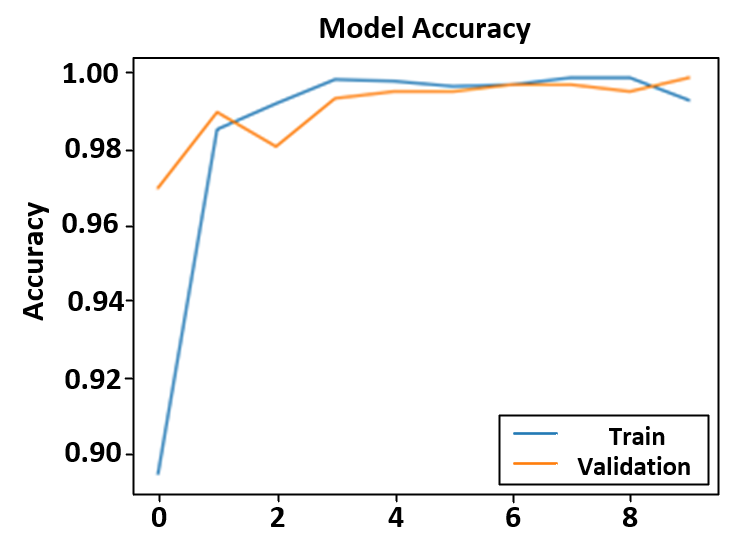 |
| **Figure 3. Confusion Matrix and Learning Curve for Classifier Applied on DCGAN-balanced Dataset.**  **(a)** Confusion matrix showing mixed classification results after DCGAN-based dataset balancing, with diagonal accuracy values: Normal (0.99), Malignant (0.75), and Benign (0.83). While benign case classification improved by 16 percentage points from baseline, malignant case accuracy declined significantly from 97% to 75%, with 24% of malignant cases misclassified as normal—a concerning outcome for clinical applications. **(b)** Training and validation accuracy curves demonstrating stable learning across 10 epochs with closely aligned performance, indicating consistent model training despite the reduced overall diagnostic reliability caused by DCGAN augmentation. | |

The drop in malignant case accuracy is especially concerning in clinical contexts, where precise detection is vital to avoid false negatives. Figure 3b shows stable learning curves, indicating that the performance issues stem from synthetic data quality rather than training instability, highlighting how generative model choice directly impacts downstream clinical utility.

**Comprehensive Performance Metrics**

The distinct performance patterns associated with each augmentation strategy are visualized in the Figure 4, providing a deeper exploration of their clinical relevance. These findings emphasize that augmentation strategies have far-reaching effects on overall diagnostic reliability. As shown in Figure 4b, the base model trained on the original imbalanced dataset (blue bars) exhibits a marked deficit in recall and F1-score for the benign class compared to the normal and malignant classes, highlighting the well-known issue of under-detection for minority classes prevalent in medical datasets. While this base approach achieved a reasonable test accuracy of 0.941 (Figure 4d), this satisfactory aggregate score belies the severe inconsistencies in per-class performance, particularly for benign lesions, which poses significant risks in clinical settings.

| (a) | (b) |
| --- | --- |
| 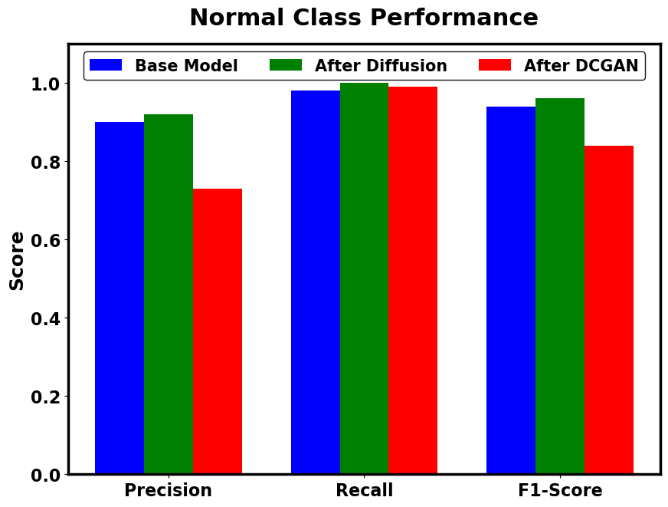 | 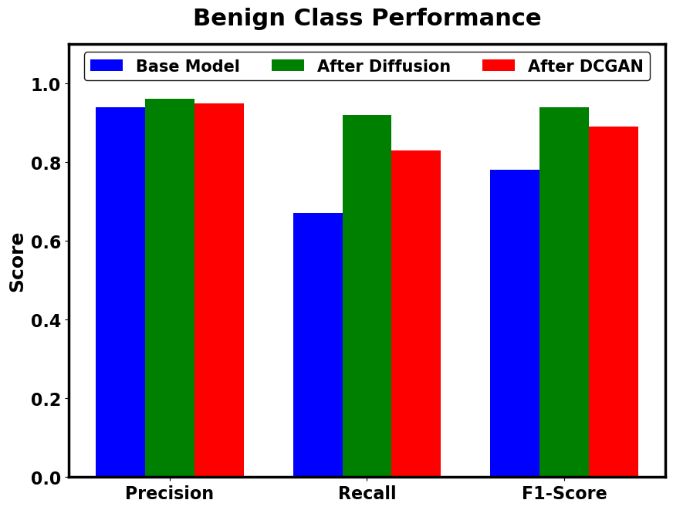 |
| (c) | (d) |
| 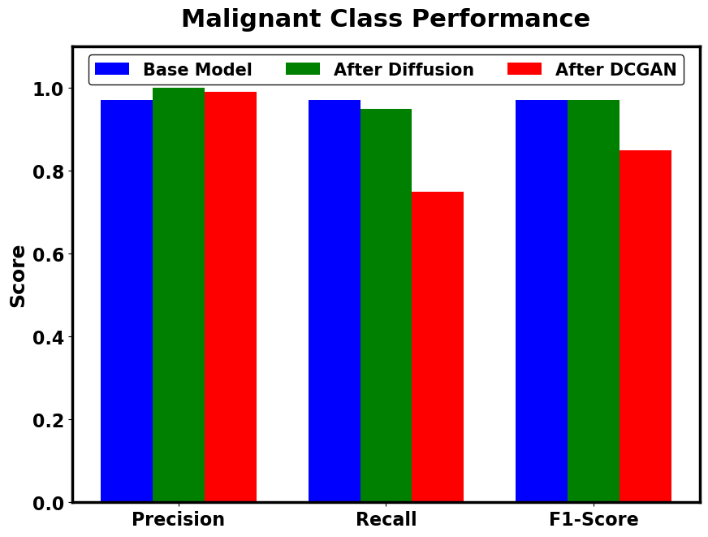 | 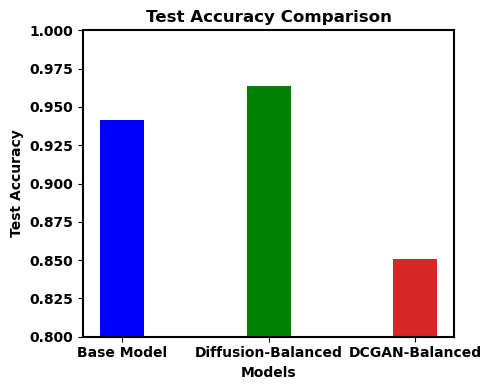 |
| **Figure 4. Class-specific and overall classification performance under different data augmentation strategies.**  (a) Normal class precision, recall, and F1-score for the base model (blue), diffusion-balanced (green), and DCGAN-balanced (red) datasets. (b) Benign class performance metrics under each augmentation methhod. (c) Malignant class performance metrics under each augmentation approach. (d) Overall test accuracy for the base model and models trained with diffusion- and DCGAN-balanced datasets. | |

Addressing these shortcomings, the diffusion model-based augmentation (green bars) led to a test accuracy increase to 0.964 (Figure 4d) and, more importantly, brought precision, recall, and F1-scores to uniformly high levels across all diagnostic classes (Figures 4a–4c). This class-level balance demonstrates that the diffusion-augmented data rectified the initial skew, allowing the model to reliably recognize normal, benign, and malignant cases—a prerequisite for trustworthy clinical implementation. In sharp contrast, the DCGAN-balanced dataset (red bars) resulted in a pronounced accuracy drop to 0.851 (Figure 4d), and Figures 4a–4c reveal that any improvement in benign case recognition was offset by substantial losses in malignant detection, raising concerns about failing to important critical pathology. These results not only highlight the necessity of high-fidelity, class-representative synthetic data but also demonstrate that not all generative augmentation approaches impart equal clinical value, with method-specific artifacts or variability potentially undermining diagnostic safety.

**Precision-Recall Analysis**

To further understand the clinical implications of the overall accuracy differences observed in Figure 8d, we examined precision-recall metrics for each diagnostic category. Precision-recall analysis in Figure 8 highlights the clinical relevance of augmentation strategies. In the diffusion-balanced dataset (green bars), precision exceeded 0.90 across all diagnostic categories, with benign class recall improving from 0.67 to 0.90 and precision holding at 0.95, raising the F1-score from 0.78 to 0.92. This balance indicates reduced false positives and enhanced true positive detection across classes. DCGAN (red bars) improved benign recall to 0.83 but led to a drop in precision for malignant (0.99→0.72) and normal (0.88→0.72) cases, reflecting inconsistencies in synthetic image quality. As shown in Figure 8d, the diffusion-balanced model achieved the highest test accuracy (0.964), outperforming both the base (0.941) and DCGAN-balanced (0.851) models, affirming the importance of evaluating generative methods based on clinical task performance rather than image fidelity alone.
