## Supplementary File S3 for "Diffusion Models vs. DCGANs for Class-Imbalanced Lung Cancer CT Classification: A Comparative Study"

**Task-Centric Evaluation Framework for Medical Image Synthesis**

**1. Overview**

This framework formalizes a task-centric evaluation protocol for validating synthetic medical images in clinical contexts. Unlike generic image quality assessment that prioritizes visual plausibility or statistical similarity, this protocol centers validation around downstream clinical task performance, with explicit safety evaluation to detect performance trade-offs that could compromise patient care.

**2. Evaluation Hierarchy**

**2.1 Primary Validation: Clinical Task Performance**

All synthetic image quality claims must be validated through performance on the intended clinical task using held-out real data. For classification tasks:

- Overall accuracy on balanced test set

- Per-class recall/sensitivity (critical for detecting false negative risks)

- Per-class precision (to assess false positive rates)

- F1-scores for balanced assessment

- Statistical validation across multiple runs (minimum n=10 with different random seeds)

**2.2 Safety Evaluation: Class-Specific Performance Analysis**

Mandatory assessment of performance across all diagnostic categories to identify unsafe trade-offs:

- Individual class recall metrics for each pathology type

- Confusion matrix analysis to detect misclassification patterns

- Statistical significance testing (paired t-tests) between augmentation strategies

- Explicit reporting of performance variance (standard deviations) across runs

- Clinical risk assessment for classes where errors carry severe consequences

**2.3 Secondary Metrics: Image Quality Assessment**

Image quality metrics serve as preliminary screening tools, not final validation:

- Multiple complementary metrics (FID, KID, IS, KL divergence) to assess different aspects

- Acknowledgment that metrics may provide conflicting rankings

- Recognition that favorable image quality scores do not guarantee clinical utility

- Use as hypothesis generators for understanding performance differences

**3. Protocol Steps**

**Step 1: Define Clinical Task and Error Consequences**

Explicitly document:

- Intended clinical application

- Relative severity of different error types (e.g., missed cancer vs. false alarm)

- Target performance thresholds for each diagnostic category

- Regulatory or clinical deployment requirements

**Step 2: Generate Synthetic Images**

- Document generative model architecture completely

- Report all training hyperparameters

- Specify number of synthetic images generated per class

- Maintain reproducibility through fixed random seeds

**Step 3: Assess Image Quality (Preliminary)**

- Evaluate using multiple complementary metrics (FID, KID, IS, KL divergence minimum)

- Calculate metrics separately for each diagnostic category

- Document any conflicting results among metrics

- Use results to identify potential issues, not as final validation

**Step 4: Downstream Task Validation (Primary)**

- Train classification model with three conditions:

a) Baseline: Original imbalanced dataset

b) Augmented: Dataset balanced with synthetic images

c) Control: Alternative augmentation strategy (if available)

- Use consistent architecture across all conditions

- Employ statistical validation (minimum 10 independent runs with different seeds)

- Report mean ± standard deviation for all metrics

**Step 5: Safety Analysis**

- Examine per-class performance for all diagnostic categories

- Calculate recall for each class (emphasis on life-threatening conditions)

- Perform paired statistical tests between augmentation strategies

- Identify any class where performance degraded compared to baseline

- Assess clinical significance of performance differences

**Step 6: Clinical Risk Assessment**

For each augmentation strategy, explicitly evaluate:

- False negative rates for critical pathologies (e.g., cancer)

- Clinical consequences of observed misclassification patterns

- Performance consistency (variance across runs)

- Whether improvements in minority classes came at expense of majority classes

**4. Reporting Requirements**

**4.1 Mandatory Reporting**

- Complete architectural details for generative models

- All image quality metric results (not just favorable ones)

- Downstream task performance with error bars (mean ± SD across runs)

- Per-class performance for all diagnostic categories

- Statistical significance tests comparing methods

- Any observed safety concerns or performance trade-offs

**4.2 Honest Interpretation**

- Acknowledge when image quality metrics conflict with task performance

- Report any classes where augmentation degraded performance

- Discuss clinical implications of performance differences

- Distinguish between statistical significance and clinical significance

**5. Application to Lung Cancer CT Classification**

Our study demonstrates this framework through comprehensive evaluation of DCGAN and Diffusion-based augmentation:

Primary Validation Results:

- Baseline: 0.9701 ± 0.0104 overall accuracy

- DCGAN-Balanced: 0.9760 ± 0.0116 overall accuracy

- Diffusion-Balanced: 0.9959 ± 0.0068 overall accuracy

Safety Analysis Revealed:

- Standard DCGAN (Supplementary File 3): Catastrophic malignant recall drop to 75%

- Improved DCGAN: Maintained malignant recall at 98.8 ± 0.9%

- Diffusion: Achieved highest malignant recall at 99.7 ± 0.8%

**Key Finding:** Image quality metrics alone (detailed in Supplementary File 1) could not predict the standard DCGAN safety failure or reliably distinguish between improved methods. Only task-based validation with safety analysis identified:

1. Life-threatening performance degradation (standard DCGAN)

2. Successful safety-preserving augmentation (improved DCGAN)

3. Superior overall performance (Diffusion)

**6. Advantages Over Image-Quality-Only Validation**

Task-centric evaluation successfully:

- Detected catastrophic failure mode (75% malignant recall) masked by competitive image quality scores

- Identified clinically meaningful performance differences not captured by image metrics

- Provided actionable guidance for method selection in clinical contexts

- Ensured augmentation strategies improve rather than compromise patient safety

Image quality metrics alone failed to:

- Predict downstream task performance reliably

- Detect unsafe trade-offs between diagnostic categories

- Distinguish between methods with similar visual quality but different clinical utility

- Account for clinical consequences of different error types

**7. Recommendations for Future Studies**

Researchers developing generative models for medical imaging should:

- Prioritize downstream clinical task validation over image quality metrics

- Always include explicit safety evaluation examining per-class performance

- Employ statistical validation across multiple runs to assess reproducibility

- Report any performance degradation in critical diagnostic categories

- Acknowledge when image quality and task performance assessments conflict

- Frame conclusions around clinical utility rather than generic image quality

**8. Conclusion**

This task-centric framework ensures that validation of synthetic medical images prioritizes patient safety and clinical utility over visual plausibility. By centering evaluation on downstream clinical performance with explicit safety analysis, this protocol prevents deployment of augmentation strategies that achieve favorable image quality metrics while introducing life-threatening performance degradations in critical diagnostic categories.
